# Simulation and experimental study on the strategy of soil auxiliary heating in a solar greenhouse in northeast China

**DOI:** 10.1101/2025.06.25.661648

**Authors:** Xin Liu, Cong Wang, Tieliang Wang, Feng Zhang, Zhanyang Xu

**Author notes:** Corresponding author (Z X).

## Abstract

When solar greenhouses in north eastern China experience extreme weather, the internal soil requires varying auxiliary heating to ensure the normal growth of crops. In this study, considering the effects of four kinds of extreme weather on solar greenhouses, four different types of heating requirements were obtained through experiences in a Liaoshen-type solar greenhouse. Four distinct heating strategies were simulated using a soil temperature prediction model. The initial temperatures for these strategies were 13°C, 11°C, 13°C, and 14°C, respectively, while the target temperature was uniformly set at 15°C. The heating durations for the respective strategies were 432,000 seconds, 432,000 seconds, 864,000 seconds, and 1,728,000 seconds. To ensure a minimum soil temperature of 15°C within the solar greenhouse, the temperature of the heating layer located 400 mm underground needed to be maintained at or above 20°C, 28°C, 18°C, and 19°C, respectively. The simulation results were validated through experimental verification, with maximum and minimum errors of 3.1% and 0.5%, respectively. Several targeted promotional suggestions were also proposed. The model exhibited high accuracy and is capable of developing optimal and precise soil heating strategies for various types of solar greenhouses in north eastern China.

## 1. Introduction

As an important tool for winter vegetable production, the solar greenhouse is widely used in the cold regions of northeast China [1]. Because of the good thermal insulation and heat storage performance of the greenhouse wall and the scientific and reasonable design of the front slope, when the outdoor temperature is above -25℃, the solar green-house can ensure the normal growth of crops without heating [2]. Solar energy accepted through the front slope is the main energy source [3]. When extreme weather occurs, such as cold waves, snow, or serial cloudy days, the climatic conditions in the greenhouse cannot adjust themselves to ensure the normal growth of crops in a short time, especially the soil temperature [4, 5]. At this time, the solar greenhouse needs auxiliary heating by reasonable methods [6].

In recent years, many researchers have studied the heating methods for solar greenhouses. Fatiha Berroug et al. [35] proposed a simple yet cost-effective rock bed storage system (RBS) for solar energy storage, which is designed to heat the air in greenhouses during night time. This system employs pebbles ranging in size from 2 to 15 centimetres to store thermal energy during the day and subsequently release it into the greenhouse at night. Hassanien et al. [7] used a kind of evacuated tube solar collector as a solar water heater assisted by an electric heat pump for greenhouse heating in Kunming, China. This heating system can increase the heated greenhouse internal air temperature by 3°C, and decrease the relative humidity by 10%. Bouadila et al. [8, 9] designed a new solar air heater by using a packed bed of spherical capsules with a latent heat storage system in Tunisia. The spherical capsules were made with phase change material (PCM) and the heat balance in different components of the heated greenhouse had been studied. At night time, the heating system can increase the greenhouse inner temperature by 5°C, and reduce relative humidity by an average of 10-20%. Xu et al. [10] proposed a water-circulating solar heat collection and release system in a greenhouse in Beijing, China. The solar heat collector was built as hollow polycarbonate sheets, and it was also used as a heating radiator for the green-house during night time. Results showed that the system’s daily average heat collection ratio was 72.1% and it can in-crease the greenhouse inner air temperature by a minimum of 3.9°C during the night time. Joudi et al. [11] made a solar air heater (SAH) system to heat an innovative greenhouse in Baghdad, Iraq. The heater was put on the roof as one structure of a greenhouse, and it can keep the inner air temperature of the greenhouse at 18°C when the SAH system covers 45% of the greenhouse roof. Zhang et al. [12] built a burning-cave hot water soil heating system in a greenhouse in Liaoning, China. The system transferred burning cave internal heat into the soil by pipes that were filled with water, and the soil temperature of the greenhouse was evenly improved by 5°C. Gourdo et al. [13] used the black plastic sleeves filled with water to create a solar heating system in a greenhouse in Agadir, Morocco. The microclimate and tomato yield in the greenhouse were studied by experiments. This simple system can improve the night time inner temperature of the greenhouse by 3.1°C, reduce the relative humidity by 10%, and increase tomato yield by 35%. In the same year, Gourdo et al. [14] also created a solar energy storing rock-bed system to heat the air temperature in a greenhouse. This system can improve the night time inner temperature of the greenhouse by 3°C and reduce the daytime inner temperature of the greenhouse by 1.9°C. Mohammadi et al. [15] carried out a new underground passive solar greenhouse to improve thermal performance, storage, and saving of heat solar energy. It provides a new idea for the greenhouse heating method.

So far, there are various methods to heat the solar greenhouse, and each heating method has a good auxiliary heating effect on the greenhouse’s inner climate. However, due to the type and location of greenhouses being different, the requirements for auxiliary heating of greenhouses are also various. Before heating a greenhouse, it is necessary to ascertain the heating requirements of greenhouses, and then generate appropriate heating strategies for different greenhouses. Numerical simulation can predict the microclimate in the solar greenhouse [16–18], and it can provide a theoretical basis for the designation of the auxiliary heating strategy in a solar greenhouse. So far, some researchers have used different ideas to predict the environmental changes in solar greenhouses. Kim et al. [19] investigated the heat flux and heat transfer in a greenhouse in South Korea by modelling method. The U-value was presented to relate to the environmental characteristics of the greenhouse. Results show that the U-value and the air temperature between the inner and exterior of the greenhouse have a highly linear relationship. Rasheed et al. [20, 21] developed a Transient Systems Simulation Program (TRNSYS) to simulate the thermal environment of different greenhouses in South Korea. The TRNSYS was used as a building energy simulation (BES) platform to make the construction strategy of greenhouses. Fatiha Berroug et al. [35] employed TRNSYS simulation and Fortran programming to predict the effects of an integrated system on drying kinetics during both diurnal and nocturnal periods. Zhang et al. [2] put forward a mathematical model to evaluate a so-lar greenhouse quantitatively. The model was developed by considering the parameters of a greenhouse, and the solar radiation allocation and spatial distribution of the greenhouse was obtained to describe the light environment quality. This model can make the optimization strategy of the greenhouse lighting regulation and planting pattern. Zhang et al. [22] developed a functional structural plant model to simulate the light and thermal performance in a Liaoshen-type solar greenhouse in China. The model includes a detailed 3D tomato canopy structure and it can be seen as the first step toward a realistic 3D greenhouse and canopy simulation. Shen et al. [23] used an energy consumption mathematical model to predict the energy consumption under different weather conditions in a Venlo greenhouse in Chongming, China. The model is established based on the principle of energy conservation and it can provide the theoretical reference for the strategy of heating in the greenhouse. Li et al. [24] built a solar heating system in a greenhouse driven by a Fresnel lens concentrator in a greenhouse in Beijing, China. The system can heat the soil directly by underground pipes and supply heat for the greenhouse in the absence of sunlight by using thermal storage in the soil. The underground soil heat transfer and heat storage performance of the greenhouse were simulated, and the strategy of pipe-buried depths can be put forward simultaneously. Zhang et al. [25] used a thermal equilibrium theory to analyse the heat transfer process of a heating system in a greenhouse in Beijing, China. The soil temperature distribution in the greenhouse was simulated by utilizing finite element method (FEM) software, and the simulation can help to make the strategy of the soil temperature control in the greenhouse. Joudi et al. [26] also developed a dynamic model to predict the innovative green-house’s inner air and soil temperature [11], and simulation results put forward the best soil sub-layer depth in the greenhouse. Jiao et al. [27] proposed a porous medium model to express the temperature and humidity characteristics of the celery crop in a double-layer film solar greenhouse in Hailiutu, China. The inner environment characteristics of the greenhouse were simulated by the computational fluid dynamics (CFD) method and the simulation can model the yields of crops in the greenhouse simultaneously. Wang et al. [28] estimated and optimized the soil thermal diffusivity of greenhouses by different methods and the optimal thermal diffusivity was used in a sinusoidal model to predict the greenhouse soil temperature at different soil depths. The results of those previous studies showed that numerical simulation could accurately predict the thermal characteristics and heat transfer in greenhouses.

Previous studies on heating in solar greenhouses have primarily focused on new methods and materials for heating, with little attention given to the heating requirements and precise control strategies tailored to different types and regions of solar greenhouses. Based on actual winter temperature and solar radiation data from the Liaoning-Shenyang region, this study uses the Liaoshen-type solar greenhouse as a case example and employs numerical simulation methods to investigate the varying heating demands and strategies for soil in solar greenhouses. Firstly, drawing on years of experimental data and using cold-resistant crops as a case study, this study investigates the various soil heating requirements for the Liaoshen-type solar greenhouses. Subsequently, numerical simulation techniques are employed to model suitable heating strategies based on the identified soil heating demands. Lastly, the accuracy of the developed model is validated through experimental verification, and recommendations for its promotion and application are pro-posed. The findings can serve as a reference for developing soil heating strategies in different types of solar greenhouses across various regions and provide a foundation for precise control methods of soil temperature in solar greenhouses.

## 2. Materials and methods

### 2.1. Experiment

The experiment was carried out in the Liaoshen-type solar greenhouse at Shenyang Agricultural University to study the variation of soil temperature in the greenhouse with the outdoor environment. The structural parameters of the greenhouse were the same as described in Zhang et al. [22]. A meteorological station (model A753, ADCON Telemetry), shown in Fig 1(a), was used to measure the climate outside the greenhouse. A temperature sensor (model TR1, AD-CON Telemetry) in the meteorological station, was used to measure the outdoor temperature of the greenhouse. The range, resolution, and accuracy of the instrument are -40°C to +60°C, 0.2°C, and <0.05°C at +20°C, respectively. A solar radiation sensor (model LP2, ADCON Telemetry) in the meteorological station, was used to measure the outdoor solar radiation of the greenhouse. The range, resolution, and accuracy of the instrument are 0 W/m2 to 2000 W/m2, 1 W/m2, and <1% at 1000 W/m2, respectively. The meteorological station was placed near the experimental greenhouse, and the temperatures and solar radiations outside the greenhouse were measured automatically. Each temperature and solar radiation data set was automatically recorded at 15-minute intervals. Soil temperature sensors (model TR2, ADCON Telemetry), shown in Fig 1(b), were used to measure the soil temperature in the greenhouse. The range, resolution, and accuracy of the instrument are - 40°C to +80°C, 0.01°C, and <0.05°C at +20°C, respectively. The soil temperature sensors were placed on the east-west axis of the greenhouse with a spacing of 1m. Each soil temperature sensor was held 200mm below the soil of the inner greenhouse, and the soil temperatures in the greenhouse were measured automatically. Each set of temperature data was automatically recorded at 15-minute intervals after calculating the average value. The soil and outdoor temperature data were collected from December to the next year March (from 2017 to 2019) continuously, and the soil heating requirements of the greenhouse can be ascertained via the measured data.

**Fig 1.**
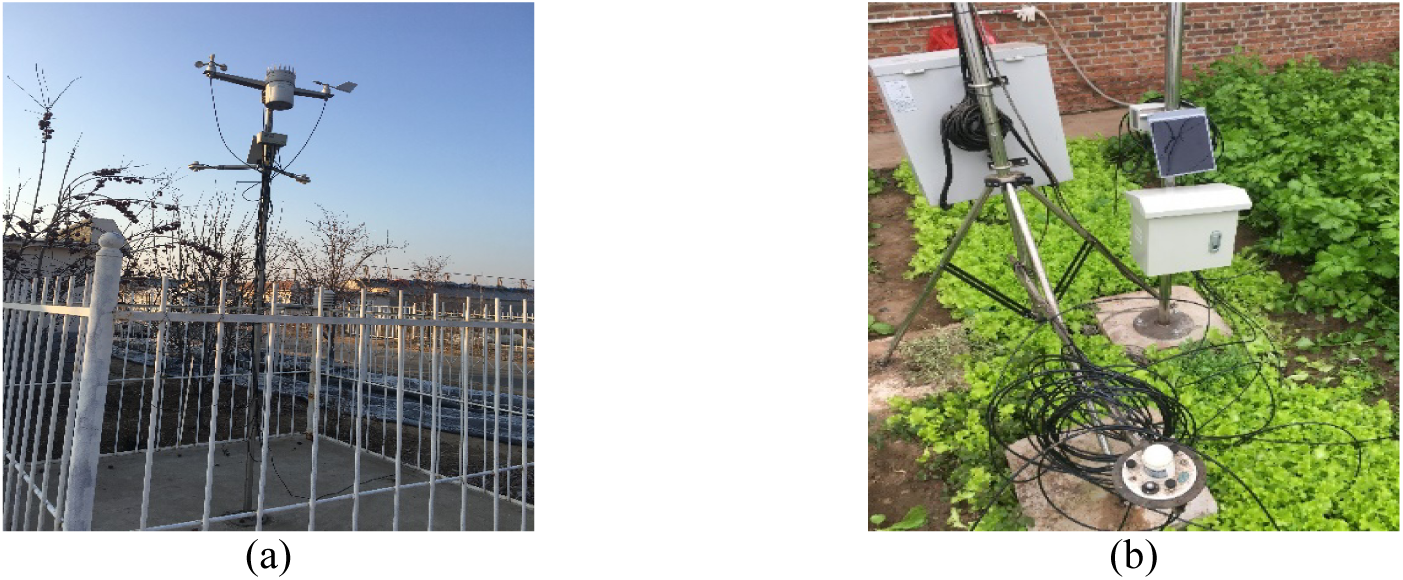
(a) Meteorological station and (b) Soil temperature sensors.

### 2.2. Simulation

In this study, commercial CFD software (Fluent, Ansys, Inc., Canonsburg, Pa.) was used to simulate the soil temperature variations in the greenhouse under heating conditions.

#### 2.2.1. Governing equations

In the greenhouse, the soil temperature at depth Z_2_ predicted from the soil temperature at depth Z_1_ can be calculated by the equation as follows [28]:

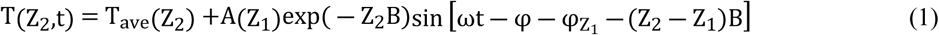

Where T(Z_2_, t) is the temperature at depth Z_2_ at any time, °C; T_ave_ (Z_2_) is the average temperature at depth Z_2_, °C; A(Z_1_) is the amplitude of the soil temperature fluctuation at depth Z_1_, °C; *φ* is the phase shift, h; ω is the radial frequency and is expressed as follows:

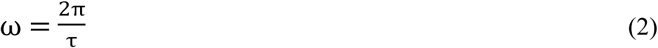

Where *τ* is the period of temperature fluctuation.

B is the inverse of the damping depth and is expressed as follows:

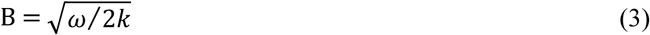

Where *k* is the soil thermal diffusivity and it can be calculated as follows [3]:

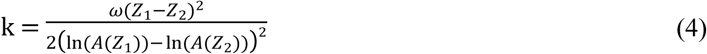

Where A(Z_1_) and A(Z_2_) are the least-squares temperature amplitudes.

#### 2.2.2. Geometric model

A 3D geometric model of the soil, shown in Fig 2(a), was built before the simulation. The dimensions of the soil model are 1m×1m×0.4m. The bottom of the geometric model is the heating plane, and it can heat the soil directly. The top of the geometric model is the surface soil in the greenhouse and the temperature can be changed via the heating plane. Other planes of the geometric model are adiabatic planes. The soil constitutes the main body of the geometric model, within which several target planes are positioned.

**Fig 2.**
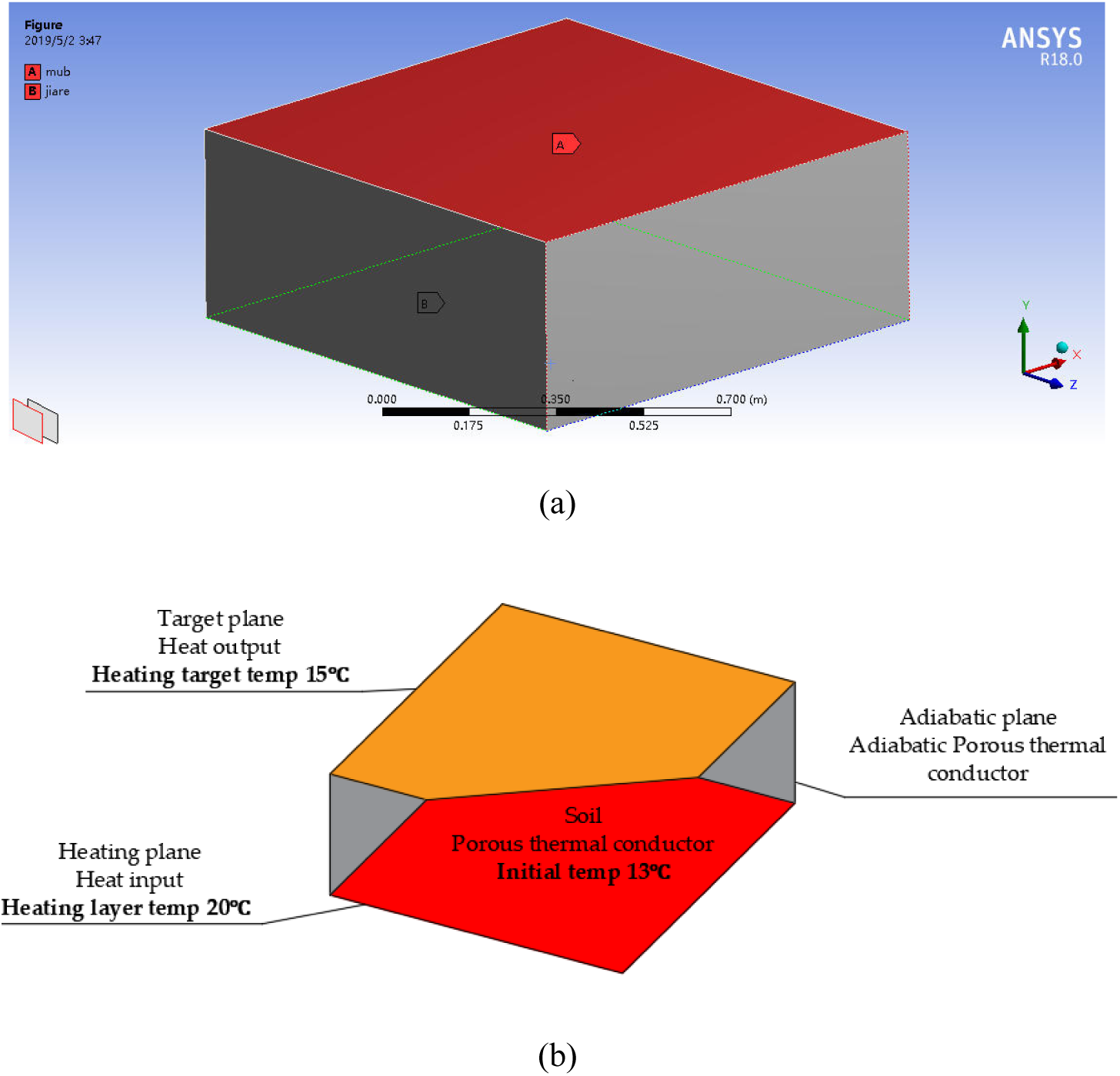
(a) Geometric model of soil and (b) Boundary condition parameters.

#### 2.2.3. Grid independence

Unstructured grid was used for the soil model in this study. Grid independence was established for the soil to get a reasonable quantity of grid cells without affecting the simulation accuracy. Three points at different locations with coordinates of (0.5, 0.2, 0.25), (0.5, 0.2, 0.5), and (0.5, 0.2, 0.75) were selected, shown in Fig 3(a), and the relationships be-tween the temperatures at these selected points and the number of grid cells were determined. The temperature variations at the three points for different quantities of grid cells were studied, as shown in Fig 3(b), and the temperatures at the selected points increased as the number of grid cells increased up to 22,680. Beyond this value, the temperatures at the selected points remained constant. Therefore, the minimum quantity of grid cells that did not affect the accuracy of the simulation was determined to be 22,680, and this value was used in this study.

**Fig 3.**
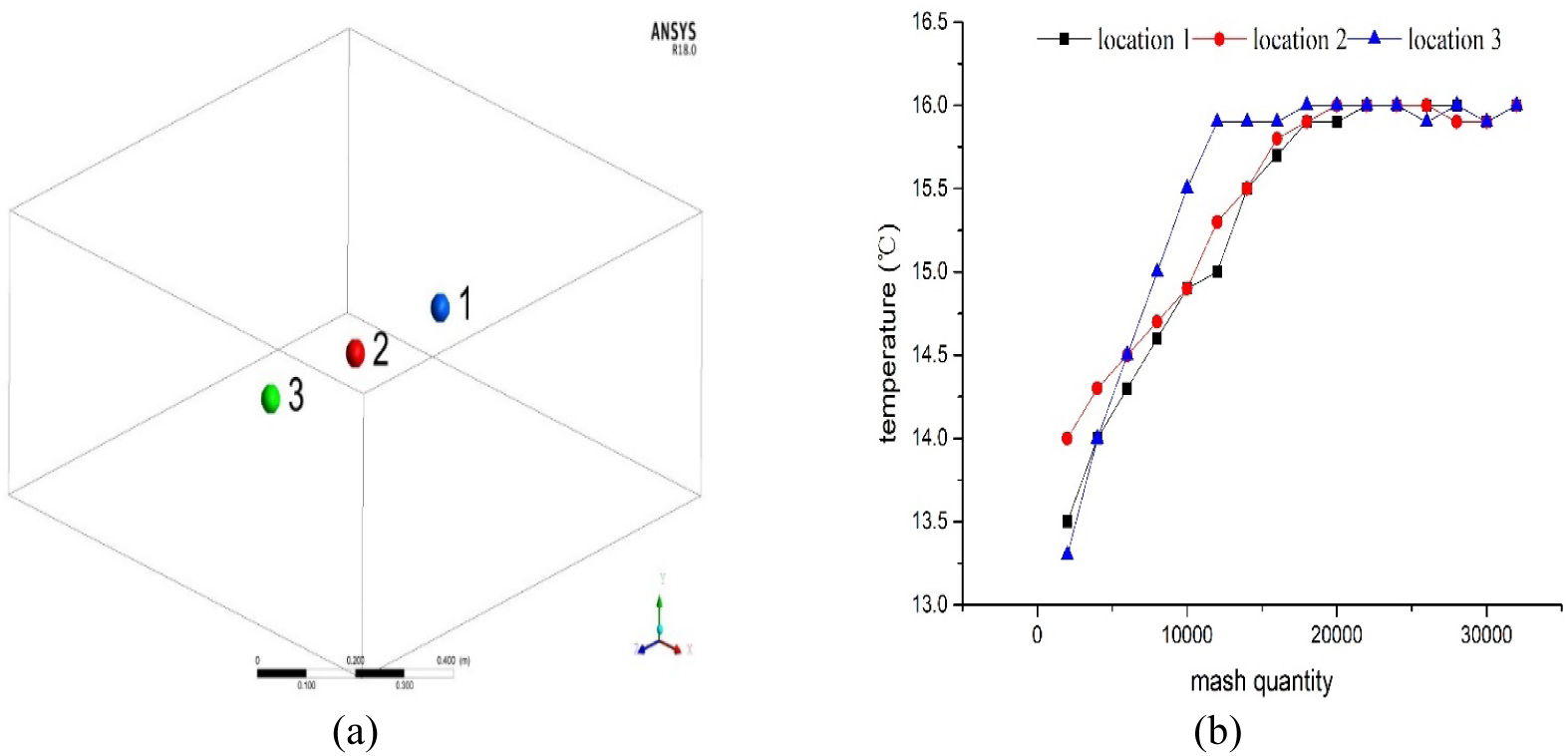
(a) Locations of three selected points and (b) Variations of temperature with mesh quantity.

#### 2.2.4. Boundary conditions

The boundaries considered in this study were the heating plane, target plane, adiabatic plane, and soil. The conditions of each boundary were the heat input, heat output, adiabatic, and porous thermal conductor, respectively. The thermos physical parameters of the porous thermal conductor (soil) were set in the simulation model. The thermal conductivity, thermal capacity, moisture content, and porosity of the soil were 1 Wm^-1^ K^-1^, 1.25×109 Jm^-3^, 25%, and 0.6, respectively. The parameters of the boundary conditions are shown in Table 1. All the boundary conditions used in this study were based on experiments. At the same time, the soil was set as homogeneous and the thermal conductivity and thermal capacity will not be changed over time. There was only one heat input at the bottom of the soil model and the heat transfer direction was vertical. The energy effects of water transport in the soil were ignored in this study. The Boundary condition parameters are shown in Fig 2(b).

**Table 1.** Boundary conditions of the soil.

| Boundary | Conditions | Moisture content | Porosity | Thermal conductivity | Thermal capacity | Temp |
| --- | --- | --- | --- | --- | --- | --- |
| Heating plane | Heat input | — | — | — | — | Changed |
| Target plane | Heat output | — | — | — | — | Changed |
| Adiabatic plane | Adiabatic | — | — | — | — | — |
| Soil | Porous thermal conductor | 25% | 0.6 | $1\text{wm}^{-1}\text{k}^{-1}$ | $1.25\times 109\text{Jm}^{-3}$ | Changed |

### 2.3. Validation

The validation experiments were conducted to verify the accuracy of the soil heating strategy made by simulation. The burning cave [30] was used to heat the soil in the greenhouse directly. Five temperature sensors were used to verify that the temperature on the roof of the burning cave matched the heating plane formulated by simulation. These sensors were placed atop the burning cave to measure the roof temperature. The locations and data collection methods of the sensors were detailed in Xu et al. [31]. The validation experiments were performed in the same kind of Liaoshen- type solar greenhouse at the same time. The sensors and data collection methods were the same as described in this study.

## 3. Results and discussion

### 3.1. The soil heating requirements of the greenhouse

The soil heating requirements are based on the types of crops in the greenhouse as well as the outdoor temperature and solar radiation. Monthly average values of outdoor temperature and solar radiation during the test period are shown in Table 2. In this study, the lowest soil temperature suitable for the growth of hardy vegetables was set as the critical temperature. When the soil temperature in the greenhouse is below 15°C, the soil in the greenhouse should be heated in time [32].

**Table 2.** Monthly average values of outdoor temperature and solar radiation.

| Month | Average temperature (°C) | Average solar radiation (Wm<sup>-2</sup>) |
| --- | --- | --- |
| November | -1.1 | 52.1 |
| December | -6.7 | 41.2 |
| January | -9.2 | 48.4 |
| February | -4.9 | 78.3 |

Fig 4 shows the variation of greenhouse interior soil temperature with outdoor temperature and solar radiation in November. As shown in Fig 4, the soil temperature in the greenhouse was affected by both outdoor temperature and solar radiation. The soil in the greenhouse did not need to be heated during most of November, and the average temperature of the soil can stay above 15°C. However, the greenhouse was hit by conditions of snowfall on November 17th and 18th. The maximum daily solar radiation was below 50 W m^-2^ during that time. Then the outdoor temperature began dropping abruptly on November 19th and remained low for three consecutive days. In the event of sudden and sharp decreases in temperature, the self-regulating capacity of solar greenhouses is relatively limited. During this period, the internal greenhouse temperature may not adequately satisfy the growth requirements of crops. The daily maximum and minimum outdoor temperatures dropped by 20°C and 15°C respectively, and the average outdoor temperature was -13°C during that time. Because of the extreme weather, the soil temperature in the greenhouse also dropped abruptly at the same time. The maximum and minimum daily soil temperatures in the greenhouse dropped by 8°C and 5°C respectively, and the average soil temperature in the greenhouse was 13°C from November 17th to 21st. This has a significant impact on the crops in the greenhouse, which may lead to reduced yields or even the death of the crops. So the soil temperature in the greenhouse needs to be raised by 2°C for 5 days.

**Fig 4.**
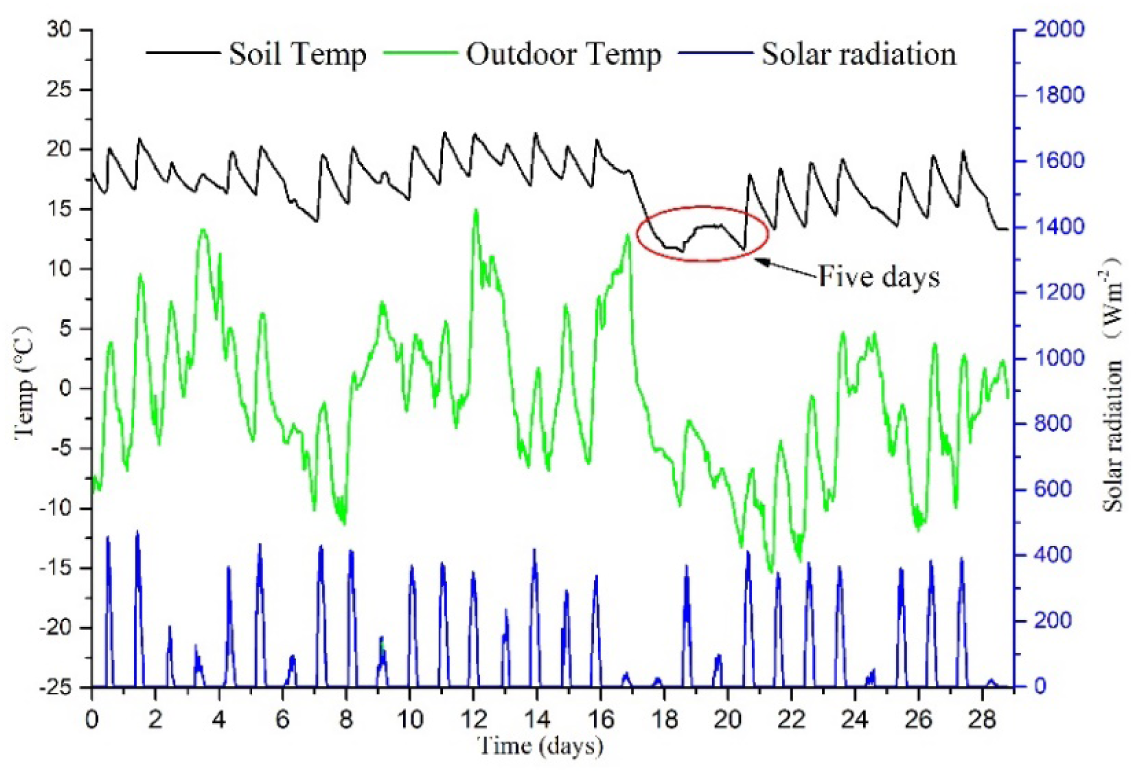
Variation of greenhouse interior soil temperature with outdoor temperature and solar radiation in November.

Fig 5 shows the variation of greenhouse interior soil temperature with outdoor temperature and solar radiation in February. As shown in Fig 5, the greenhouse was hit by conditions of overcast weather and low-temperature weather simultaneously from February 16^th^ to 24^th^. There were 4 days of overcast weather, and the maximum daily solar radiation was below 50 W m^-2^ each day. There were 5 days of low-temperature weather, and the average outdoor temperature was -13°C during that time. Under the combined influence of persistent low temperatures and prolonged over-cast conditions, the solar greenhouse experiences restricted access to external energy sources. Upon depletion of the internally stored energy, the ground temperature within the greenhouse is also likely to undergo a sharp decline. The maximum soil temperature in the greenhouse dropped by 6°C on February 16th and the soil temperature was not in-creased significantly in the next 9 days. In this scenario, it is imperative to provide heating for the greenhouse. The average soil temperature in the greenhouse was 13°C from February 16^th^ to 25^th^, and the soil temperature in the greenhouse needs to be raised by 2°C for 10 days.

**Fig 5.**
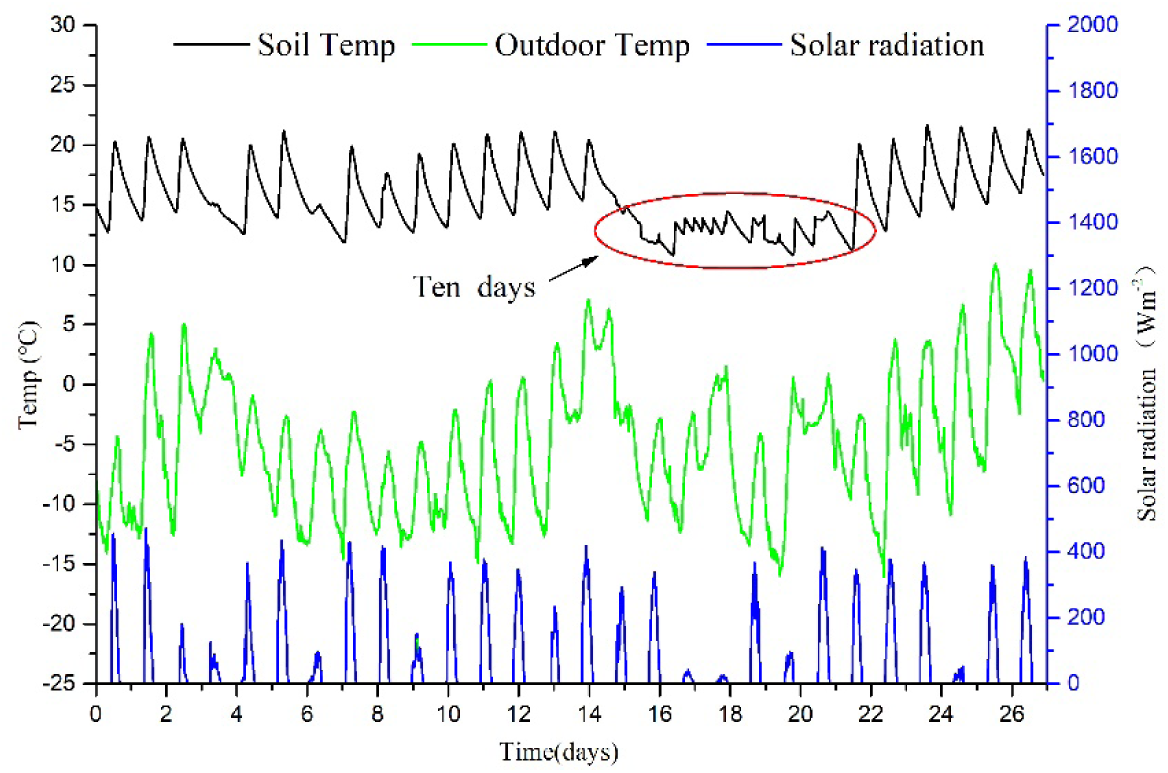
Variation of greenhouse interior soil temperature with outdoor temperature and solar radiation in February.

Fig 6 shows the variation of greenhouse interior soil temperature with outdoor temperature and solar radiation in January. As shown in Fig 6, the greenhouse was hit by the condition of low-temperature weather on January 7^th^. The outdoor maximum temperature dropped 8°C on that day, and the outdoor temperature continued to drop over the next 7 days. The maximum and minimum outdoor temperatures were -10°C and -22°C respectively. Even under optimal lighting conditions, if a solar greenhouse experiences an extended period of low temperatures, the soil temperature within the greenhouse will remain consistently low. The average soil temperature in the greenhouse was 13°C from January 7^th^ to 16^th^, and the soil temperature in the greenhouse needs to be raised by 2°C for 10 days. The greenhouse was also hit by conditions of snowy weather and low-temperature weather simultaneously from January 24^th^ to 28^th^. The maximum daily solar radiation was below 50 W m^-2^ each day and the average outdoor temperature was -13°C during that day. Under the cumulative impact of multiple extreme weather events, the soil temperature within solar greenhouses is likely to experience a sharp decline and remain at a low level for an extended period. At this point, the solar greenhouse requires sustained and efficient rapid heating. The average soil temperature in the greenhouse was 11°C from January 24^th^ to 28^th^, and the soil temperature in the greenhouse needs to be raised by 4°C for 5 days.

**Fig 6.**
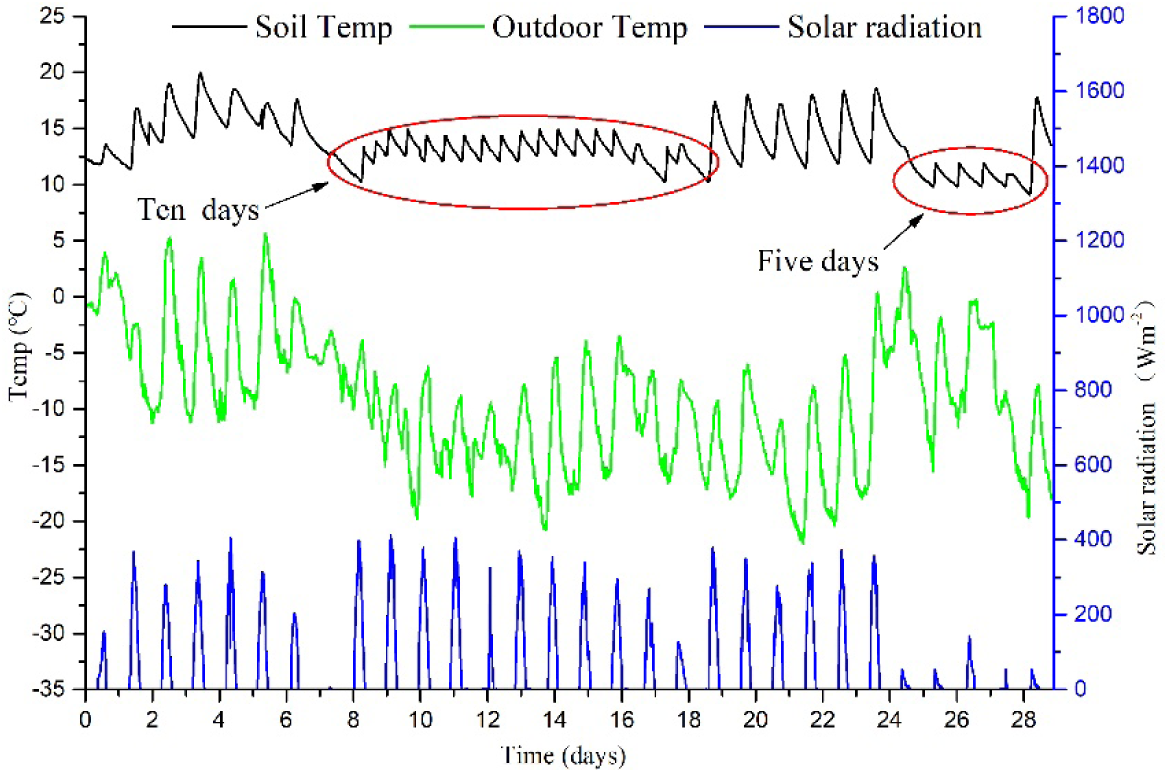
Variation of greenhouse interior soil temperature with outdoor temperature and solar radiation in January.

Fig 7 shows the variation of greenhouse interior soil temperature with outdoor temperature and solar radiation in December. As shown in Fig 7, the outdoor temperature continued to drop from December 6^th^ to 14^th^, the minimum outdoor temperature dropped from -8°C to -22°C. Then the outdoor temperature began to rise gradually from December 15^th^ to 25^th^, and the minimum outdoor temperature rose from -22°C to -3°C. However, the greenhouse was hit by overcast weather for 3 days from December 15^th^ to 25^th^, and the maximum daily solar radiation was below 40 Wm^-2^ each day. The soil maximum temperature and minimum temperature in the greenhouse fluctuated from around 15°C from December 6^th^ to 25^th^. This situation demonstrates the inherent temperature regulation capability of solar greenhouses; however, the control capacity is inherently constrained. And the soli minimum temperature was below 15°C for most of that time. The average soil temperature in the greenhouse was 14°C from December 6^th^ to 25^th^, and the soil temperature in the greenhouse needs to be raised by 1°C for 20 days.

**Fig 7.**
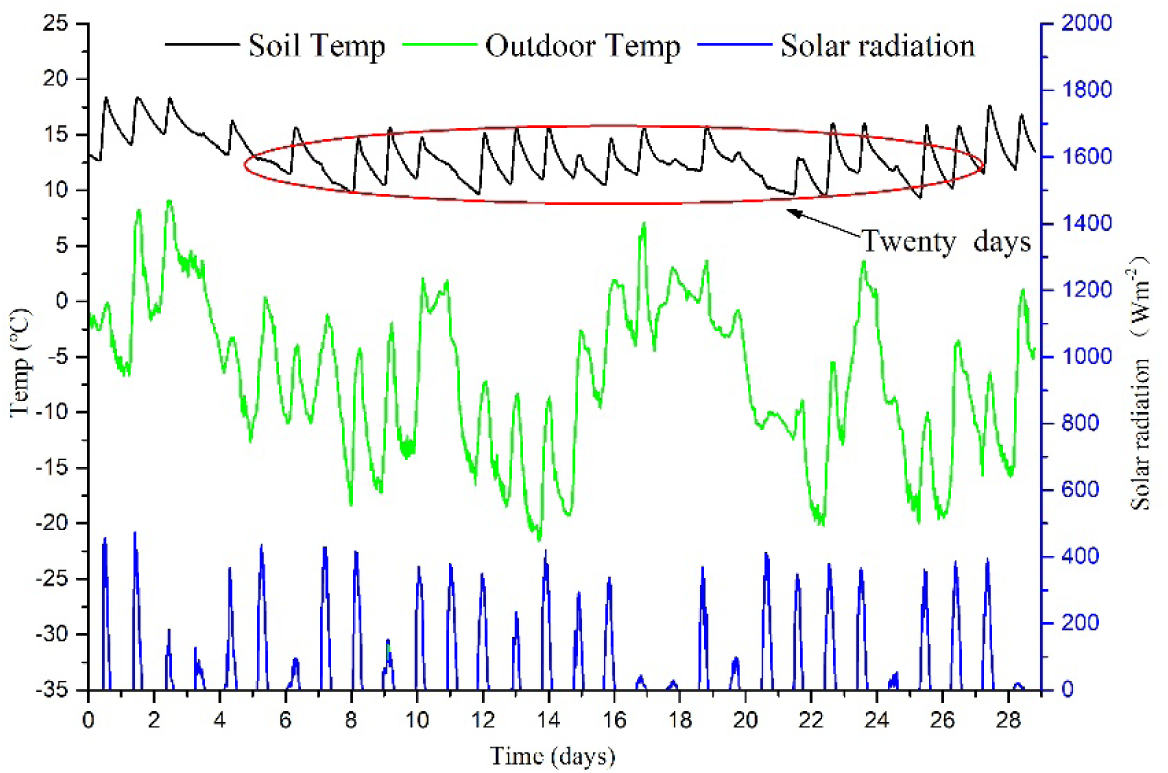
Variation of greenhouse interior soil temperature with outdoor temperature and solar radiation in December.

According to the experimental results, four types of auxiliary heating are required in Liaoshen- type solar greenhouses in Northeast China. Type A, when the outdoor temperature is at a high level and there is snowfall or a sudden drop in temperature at the same time, the average soil temperature in the greenhouse should be raised by at least 2℃ for 5 days. Type B, when the outdoor temperature is at a low level and there is snowfall or a sudden drop in temperature at the same time, the average soil temperature in the greenhouse should be raised by at least 4℃ for 5 days. Type C, when the outdoor temperature is in a state of continuous low temperature, the average soil temperature in the greenhouse should be raised by at least 2℃ for 10 days. Type D, when the greenhouse is hit by overcast weather or brief cooling weather, the average soil temperature in the greenhouse should be raised by at least 1℃ for 20 days. The auxiliary heating types are shown in Table 3.

**Table 3.** Auxiliary heating types for the greenhouse.

| Heating types | Heating time duration (days) | Temperature increased (°C) | Adverse weather description |
| --- | --- | --- | --- |
| Type A | 5 | 2°C | Snow and low temperature |
| Type B | 5 | 4°C | Snow and continuous low temperature |
| Type C | 10 | 2°C | Continuous low temperature |
| Type D | 20 | 1°C | Continuous low temperature or overcast weather |

### 3.2. The heating strategy of the greenhouse

Based on experimental data, the heating strategy of the soil in the greenhouse was determined by simulation. Under heating type A, the initial temperature of the soil is 13 ℃, and the target temperature is 15℃. The soil horizontal plane temperature distribution at 200mm underground is shown in Fig 8(a), and the soil vertical temperature distribution is shown in Fig 8(b). In Fig 8, the distribution of soil temperature at a depth of 200mm below the ground shows a symmetrical pattern, with the highest temperature (16℃) in the central region. The temperature gradually decreases from the center towards the surrounding areas. The boundary region of the model is defined by the lowest temperature, which is initially set at 13℃. The temperature distribution in the vertical plane of the soil is axisymmetric, with the highest temperature at the bottom of the soil (20℃) and the lowest temperature around the soil (13℃). Due to the presence of a heating layer located 400mm underground, the temperature of the soil increases horizontally as the depth of the soil increases. For heating type A, it is essential to maintain the temperature of the heating layer at a minimum of 20°C. Additionally, the mean temperature of the soil at a depth of 200mm underground can exceed 15°C in most areas, meeting the temperature requirements for the suitable growth of hardy and semi-hardy vegetables.

**Fig 8.**
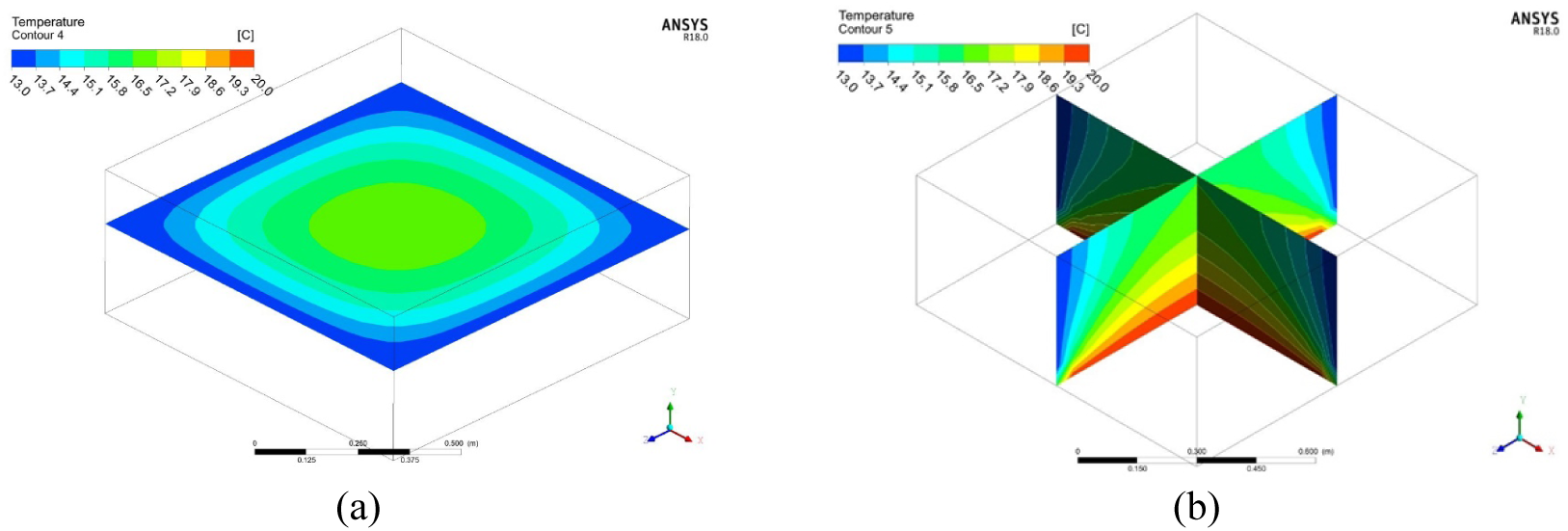
Soil temperature distribution on the horizontal plane (a) and vertical plane (b) under type A heating.

Under heating type B, the initial temperature of the soil is 11 ℃, and the target temperature is 15℃. The soil horizontal plane temperature distribution at 200mm underground is shown in Fig 9(a), and the soil vertical temperature distribution is shown in Fig 9(b). In Fig 9, the distribution of soil temperature at a depth of 200mm below the ground also shows a symmetrical pattern, but with significant temperature differences. The central region experiences the highest temperature, reaching 17.5℃ due to faster heating, while the temperature gradually decreases towards the surrounding regions. The lowest temperature occurs at the boundary region of the model, with the initial soil temperature being 11℃. The temperature distribution in the vertical plane of the soil is axisymmetric, with the highest temperature at the heating layer (28°C) and the lowest temperature near the initial soil temperature (11°C). For heating type B, it is essential to maintain the temperature of the heating layer at a minimum of 28°C.Meanwhile, the mean temperature of the soil at a depth of 200mm underground can exceed 15.5°C in most areas.

**Fig 9.**
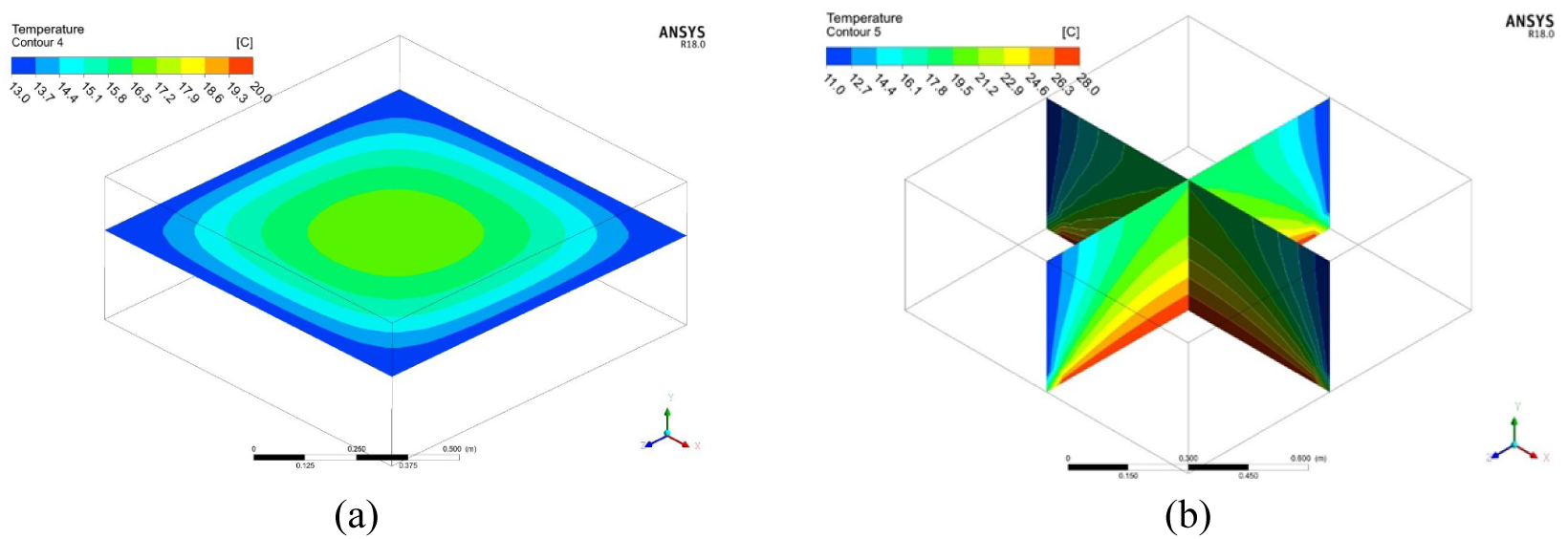
Soil temperature distribution on the horizontal plane (a) and vertical plane (b) under type B heating.

Under heating type C, the initial temperature of the soil is 13 ℃, and the target temperature is 15℃. The soil horizontal plane temperature distribution at 200mm underground is shown in Fig 10(a), and the soil vertical temperature distribution is shown in Fig 10(b). In Fig 10, after being heated for a while, the soil has a small temperature difference. The highest temperature in the central region is 16.5℃, gradually decreasing towards the surrounding areas, with the lowest temperature being 15℃ at the boundary. The temperature distribution in the vertical plane of the soil is axisymmetric, with the highest temperature of 18°C at the bottom and the lowest temperature elsewhere. Over time, the long-term heating method has raised the lowest temperature in the soil to the target temperature of 15°C. For heating type C, when the temperature of the heating surface is 18°C, the temperature of all areas of the underground 200mm soil can reach more than 15°C. For heating type C, when the temperature of the heating layer is 18°C, the temperature of all areas of the underground 200mm soil can reach more than 15°C.

**Fig 10.**
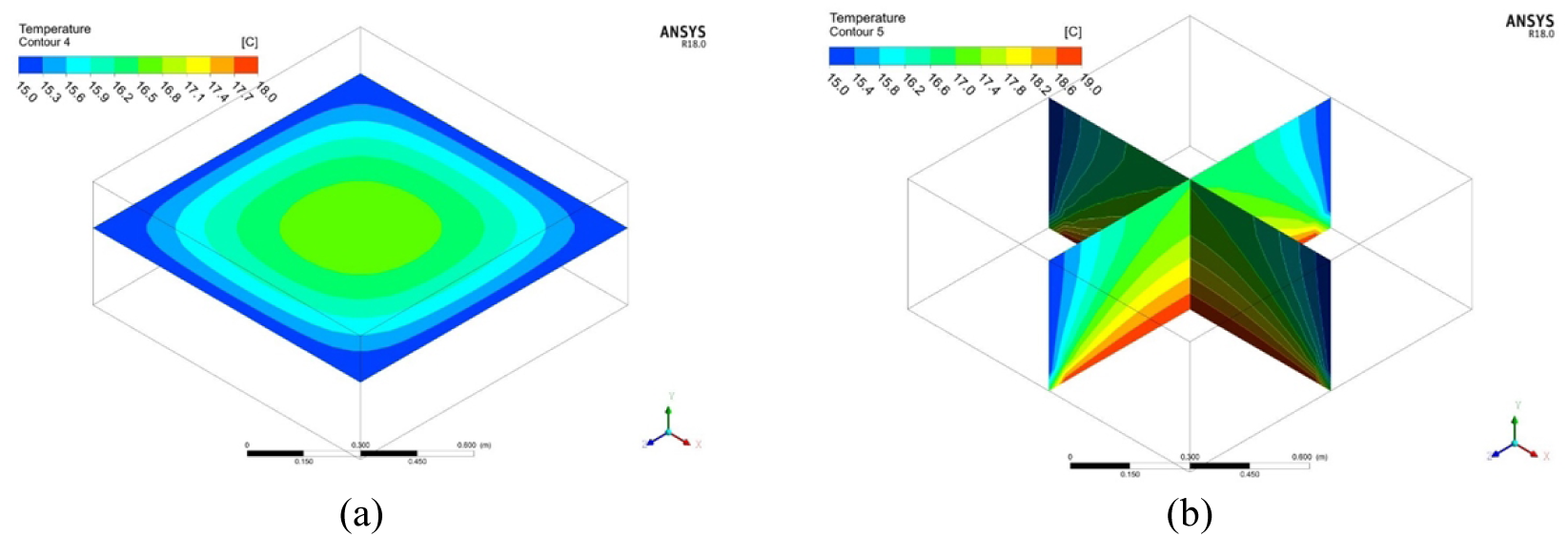
Soil temperature distribution on the horizontal plane (a) and vertical plane (b) under type C heating.

Under heating type D, the initial temperature of the soil is 14 ℃, and the target temperature is 15℃. The soil horizontal plane temperature distribution at 200mm underground is shown in Fig 11(a), and the soil vertical temperature distribution is shown in Fig 11(b). In Fig 11, the distribution of soil temperature in the horizontal plane and the vertical plane is similar to that of type C. After prolonged heating, the highest temperature is 16℃ at the center of the soil at a depth of 200mm underground, while the lowest temperature of 15℃ is at the boundary area of the model. The highest bottom temperature of the soil vertical plane is 19℃, with the surrounding temperature being the lowest, but the lowest temperature in the soil reaches the target temperature of 15℃. For heating type D, the heating time is longer, requiring the heating layer to reach a temperature of 19°C, enabling all areas of the underground 200mm soil to reach over 15°C.

**Fig 11.**
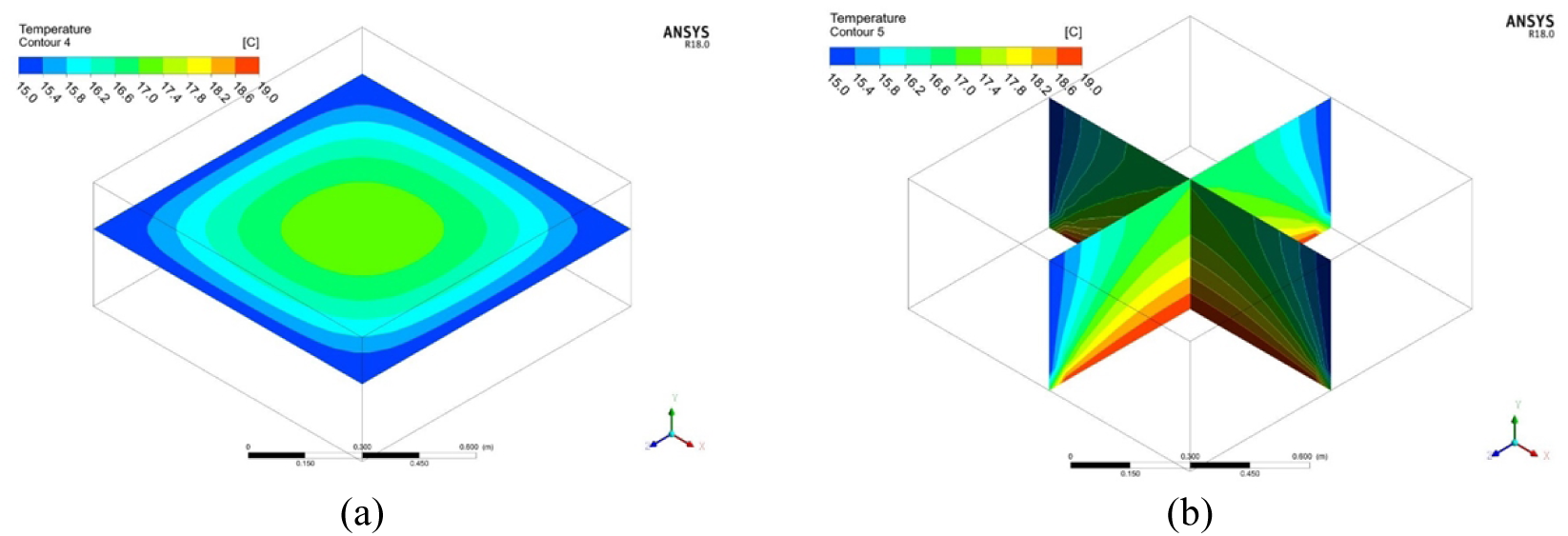
Soil temperature distribution on the horizontal plane (a) and vertical plane (b) under type D heating.

The soil is heated differently based on its initial temperature and the duration of heating. Short-term rapid heating (type B) and long-term heating (type D) require higher temperatures in the heating layer, with temperatures of 28°C and 19°C, respectively. When the initial temperature and the target temperature of the soil are the same, the temperature of the heating layer is higher for short-term heating (type A) than for long-term heating (type C). The temperature of the heating layer for type A is 20°C, while for type C it is 18°C. The heating strategy of the greenhouse is shown in Table 4. In practical greenhouse production, the temperature of the heating layer not only fluctuates in response to variations in the outdoor climate but also aligns with the specific requirements of different crops. For example, when cultivating warm-loving crops in the greenhouse, it is necessary to maintain a higher temperature in the heating layer over an extended period.

**Table 4.** The heating strategy of the greenhouse.

| Heating strategy | Soil initial temp (°C) | Soil target temp (°C) | Heating duration (s) | Heating layer temp (°C) |
| --- | --- | --- | --- | --- |
| Type A | 13 | 15 | 432,000 | 20 |
| Type B | 11 | 15 | 432,000 | 28 |
| Type C | 13 | 15 | 864,000 | 18 |
| Type D | 14 | 15 | 1,728,000 | 19 |

### 3.3. Results of validation experiments

The initial 50,000 seconds of experimental data from four heating strategies were selected for validation. Three temperature measuring points in the soil were selected to ensure the accuracy of the model. The average temperature value from these three points was then compared with the simulated data. The coordinates of the temperature measuring points can be found in Fig 3(a). The error diagram comparing four types of heating strategies between soil temperature simulation and experimental results is shown in Fig 12. As shown in Fig 12, the soil temperature variation trend was similar across the four heating strategies, with a faster rate of warming observed when the heating layer temperature was higher. After 40000 seconds of heating, each heating strategy stabilizes, and the soil temperature reaches the target of over 15°C. When comparing the experimental and simulated data, it is evident that the simulated values have a large error in the initial heating stage. When the soil temperature rises quickly, heat will be transferred to the greenhouse through the soil surface. However, this heat transfer process is not accounted for in the simulation, resulting in an overestimation of simulated soil temperatures. The maximum error is 3.1%, occurring at 10000 seconds for type B. When each heating strategy stabilizes in a stable state, the heat stays in the soil, and the numerical simulation error decreases, with the error value being less than 1.5%. The minimum error reached 0.5%, occurring at 45000 seconds for type C. The model has high accuracy, allowing for precise formulation of the soil heating strategy.

**Fig 12.**
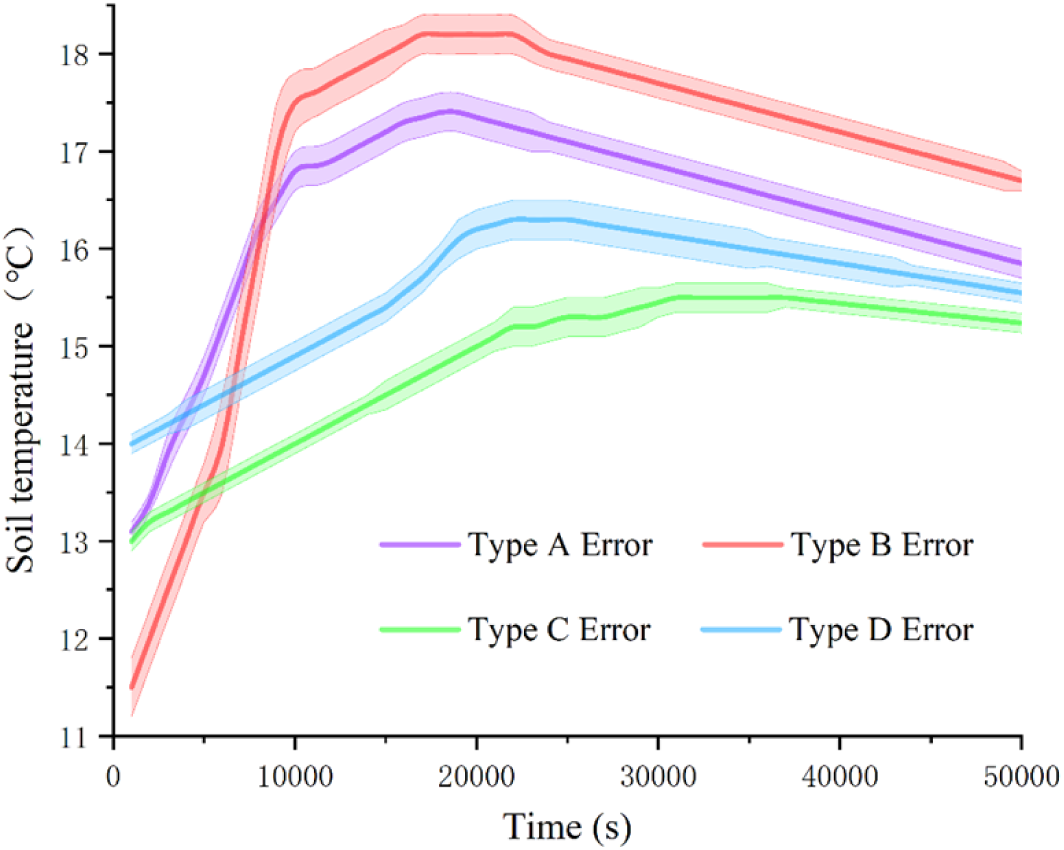
Error diagram comparing four types of heating strategies between soil temperature simulation and experimental results.

### 3.4. Discussion

The soil thermal diffusivity *κ*, a critical parameter governing the temporal and spatial dynamics of soil temperature, exhibits significant variation depending on factors such as soil texture, composition, and environmental conditions. Different *κ* correction methods should be adopted for different soil types to improve the accuracy of soil temperature simulation.

In sandy soils, which are characterized by coarse particles and high porosity, thermal conductivity (*λ*) increases linearly with moisture content (*θ*) due to the formation of enhanced antiparticle water bridges. Simultaneously, volumetric heat capacity (*ρ*C) rises proportionally with *θ*. In contrast, clay soils, known for their fine-grained structure and relatively higher thermal conductivity in the dry state, exhibit a more muted response to changes in moisture content. Organic-rich soils introduce additional complexity to this relationship, as the presence of low-density organic matter tends to reduce *λ* while increasing *ρ*C. This leads to a significant reduction in thermal diffusivity (*κ*) compared to mineral-dominated soils, which is a critically important factor for soil in solar greenhouses operating under extremely cold conditions. Environmental dynamics, particularly freeze-thaw cycles, introduce significant nonlinearities into the thermal conductivity (*κ*). The phase transition of pore water modifies both thermal diffusivity (*λ*) and volumetric heat capacity (*ρ*C). Ice formation notably enhances thermal diffusivity (*λ*) due to the higher thermal conductivity of ice compared to liquid water, while simultaneously increasing volumetric heat capacity (*ρ*C) as a result of latent heat effects. Consequently, this leads to a moderate net increase in thermal conductivity (*κ*), highlighting the necessity of coupled thermal-hydrological models for accurately capturing transient phase transitions. If layered soil systems are present in greenhouse soil, it is necessary to achieve vertical integration of *κ* using harmonic averaging (*κ_eff_*), which imposes a more severe penalty on low-*κ* layers. This phenomenon has been observed in stratified sandy-clay systems, where *κ_eff_* converges toward the lower κ value of the clay sublayer.

Future research should prioritize multi-scale validation by integrating remote sensing and machine learning techniques [33,34] to address spatial heterogeneities, especially in organic-rich or cryogenic soils where legacy models frequently exhibit limitations. The synergy between process-based modelling and data-driven approaches will play a critical role in enhancing predictive capabilities across agro ecology, climate science, and geotechnical engineering.

## 4. Conclusions

This study determined the heating demand for solar greenhouse soil in northeast China through experimentation. Subsequently, the heating strategies based on this greenhouse soil were devised using numerical simulation. Addition-ally, the effectiveness of the numerical simulation heating strategy was verified through experimentation. Some important findings and conclusions are summarized below:

(1) The Liaoshen-type solar greenhouses in northeast China exposed to different types of extreme weather require four types of soil-assisted heating. When a greenhouse encounters snow and low-temperature weather, Type A heating is required. If the greenhouse experiences snow along with prolonged low temperatures, Type B heating should be used. In cases of continuous low-temperature conditions, Type C heating is necessary. Lastly, when the greenhouse faces either persistent low temperatures or overcast weather, Type D heating is needed. Type A requires raising the soil temperature by at least 2°C for 5 days. Type B requires raising the soil temperature by at least 4°C for 5 days. Type C requires raising the soil temperature by at least 2°C for 10 days. Type D requires raising the soil temperature by at least 1°C for 20 days.

(2) Based on the four types of soil-assisted heating, it is necessary to position a heating layer 400mm below the soil in the greenhouse to heat the soil. Corresponding to heating strategies A, B, C, and D, the temperature of the heating layer should be at least 20°C, 28°C, 18°C, and 19°C, respectively. For short-term rapid heating (Type B) and long-term heating (Type D), due to the significant difference between the initial soil temperature and the target temperature, the temperature of the heating layer should be set higher than that of other conditions. When the initial soil temperature and the target temperature are identical, the temperature of the heating layer is higher for short-term heating (Type A) compared to long-term heating (Type C). The simulation results indicate that the temperature of the soil in the greenhouse can exceed 15 °C after being heated.

(3) The experimental verification confirmed that all four simulated heating strategies demonstrated effective heating, and the soil temperature after heating by the burning cave can reach over 15°C. The model demonstrates its accuracy with maximum and minimum error values of 3.1% and 0.5% for simulation and experiment, respectively. In model calculations, accounting for the influence of different soil types on soil thermal diffusivity can enhance the accuracy of numerical simulations. It can develop precise soil heating strategies for various solar greenhouses in northeast China.

## Author Contributions

Conceptualization: T.W. and Z.X.; Data curation: X.L., Z.X., and F.Z.; Model analysis: C.W. and Z.X.; Software: F.Z. and C.W.; Validation: X.L. and F.Z.; Writing–original draft: X.L. Writing– review & editing: Z.X.; All authors have read and agreed to the published version of the manuscript.

